# Nanoparticle fragmentation at solid state under single picosecond laser pulse stimulation

**DOI:** 10.1101/2021.06.02.446841

**Authors:** Peiyuan Kang, Yang Wang, Blake A. Wilson, Jaona Randrianalisoa, Zhenpeng Qin

## Abstract

Understanding the laser-nanomaterials interaction including nanomaterial fragmentation has important implications in nanoparticle manufacturing, energy, and biomedical sciences. So far, three mechanisms of laser-induced fragmentation have been recognized including non-thermal processes and thermomechanical force under femtosecond pulses, and the phase transitions under nanosecond pulses. Here we show that single picosecond (ps) laser pulse stimulation leads to anomalous fragmentation of gold nanoparticles that deviates from these three mechanisms. The ps laser fragmentation was weakly dependent on particle size, and it resulted in a bimodal size distribution. Importantly, ps laser stimulation fragmented particles below the melting point and below the threshold for non-thermal mechanism. This study reveals a previously unknown regime of nanoparticle fragmentation.

---

The interaction between light and nanomaterials leads to many interesting phenomena, such as near-field enhancement^1^, surface-enhanced Raman scattering^2^, and photothermal effects^3^. These phenomena have been harnessed for a variety of applications. For example, near-field enhancement can be used for optical trapping of molecules^4^, while surface-enhanced Raman scattering and photothermal effects have found applications in imaging^5, 6^, sensing^7, 8, 9, 10^ and actuation of biological activities^11, 12^. However, nanoparticles subjected to pulsed laser irradiation may also undergo laser-induced fragmentation, in which the original nanoparticles are fragmented into smaller particles^13^. The previously noted applications often rely on maintaining the plasmonic nanomaterial’s structural integrity during laser irradiation, as any alterations to nanoparticle morphology may adversely affect functionality^14, 15^. Moreover, laser-induced fragmentation has been used as an alternative form of green manufacturing for difficult-to-synthesize nanomaterials^16, 17, 18, 19, 20, 21^, an application which also requires accurate control of the structural integrity of plasmonic nanostructures under laser irradiation. It is therefore critical to understand what conditions lead to laser-induced fragmentation and what fundamental mechanisms control the fragmentation.

Three major categories of mechanisms have been proposed for laser-induced nanoparticle fragmentation: i) phase transition (PT) mechanisms, ii) a thermomechanical mechanism and iii) non-thermal processes. First, according to PT mechanisms, the strong heating effect of laser causes the nanoparticle to undergo phase transitions. The fragmentation can be driven by the material phase change when the particle reaches the melting point, known as the heating-melting-evaporation route^22^. The material can also be heated to the boiling point within a short period which leads to a more violent fragmentation, known as phase explosion^22, 23, 24^. Second, in the thermomechanical mechanism, the thermomechanical force plays an important role when heating happens much faster than the mechanical relaxation time (∼3 ps for a AuNP with a diameter of 10 nm), a condition known as mechanical confinement^25, 26^. The particle may fragment when the thermomechanical force exceeds the physical strength of the material without a necessary phase transition^27^. Finally, the non-thermal processes are related to the excitation of electrons, including near-field ablation and Coulomb instability (CI). The strong near-field enhancement that occurs during femtosecond laser irradiation induces nanoparticle fragmentation without melting the material^18, 28, 29, 30^. For CI, the laser irradiation ionizes the nanoparticles and causes them to fragment when the Coulombic repulsion forces within the particles exceed the surface tension of the nanoparticle. Since the melted nanoparticles have a smaller surface tension than solid ones^31, 32^, CI tends to fragment molten particles rather than solid particles^23^. So far, most research has relied on these mechanisms to rationalize nanoparticle fragmentation^33, 34^.

Although non-thermal processes and thermomechanical mechanism have proven useful in interpreting fragmentation under femtosecond pulse laser irradiation^29, 35^, and PT is thought to be the principal mechanism of fragmentation under nanosecond (ns) pulse laser irradiation^36^, the mechanism of fragmentation under intermediate picosecond (ps) pulse laser irradiation is unclear^22, 33, 37^. Previous work has suggested that both CI and near-field ablation may be possible under ps pulse laser irradiation^23, 37, 38^, but distinguishing the two mechanisms is difficult. It is also proposed that the ps laser can trigger fragmentation with CI or PT mechanisms depending on the laser fluence^23^. Recently, it has been reported that ps laser-induced fragmentation of 54 nm gold nanoparticles is a single-step, instantaneous reaction near the gold particle boiling point^33^. However, our understanding of nanoparticle fragmentation under this regime is still largely limited.

Here, we report that stimulation with a single 28 ps laser pulse leads to anomalous gold nanoparticle (AuNP) fragmentation at temperatures below the melting transition and below the threshold for CI. We compared the ps laser-induced fragmentation to that by ns laser pulse stimulation and found that the laser fluence threshold for fragmentation under ps laser pulse stimulation was about 1-2 orders of magnitude lower than that of AuNPs subjected to ns laser pulse stimulation. Importantly, the fluence threshold for ps laser pulse fragmentation shows a weak dependence on the particle size and it emits smaller nanoparticles with distinct size distributions, in contrast to the strong size-dependence and formation of continuous extrusions by AuNPs fragmented by ns pulsed laser stimulation. Next, by analyzing the temperature changes and particle instability, we determined that fragmentation of AuNPs by ps laser pulse stimulation occurred when the AuNPs were still in the solid phase and that the fluence level was much lower than that needed to induce fragmentation by Coulomb instability. Finally, we analyzed the thermomechanical stress within the AuNPs under the ps laser irradiation using MD simulation. We found that the thermomechanical stress can only fragment molten particles. This study reveals a previously unknown regime of nanoparticle fragmentation.

## Results and Discussion

### Picosecond laser pulse induced fragmentation is weakly dependent of nanoparticle size

We first developed a simple but sensitive method to monitor nanoparticle fragmentation by extinction spectral analysis and validated the method with electron microscopy. The extinction spectrum changes of AuNP colloidal solution after laser originated from nanoparticle fragmentation, and increases with higher laser fluence (Figure 1A, steps 1&2). A 2D ratio map was used to find the extinction ratio that is sensitive to the fragmentation (Figure 1A, step 3). For instance, for 15 nm AuNP under ps laser (532 nm, full width half maximum (FWHM) = 28 ps), the fluence dependence of two representative extinction ratios is shown in Figure 1A (step 4, 500/525 nm, and 550/570 nm). The TEM images support the extinction analysis that fragmentation under ps laser occurs around 1 mJ/cm^2^ (Figure 1B). On the other hand, fragmentation under ns laser requires much higher fluence (532 nm, 300 ∼ 400 mJ/cm^2^, FWHM = 6 ns, Figure 1C), as confirmed by TEM imaging. Importantly, instead of forming small spherical fragments under ps laser, ns laser-induced fragmentation forms continuous extrusions that attach onto original particles. These results confirm that extinction analysis is a feasible method to monitor nanoparticle fragmentation under pulsed lasers.

**Figure 1.**
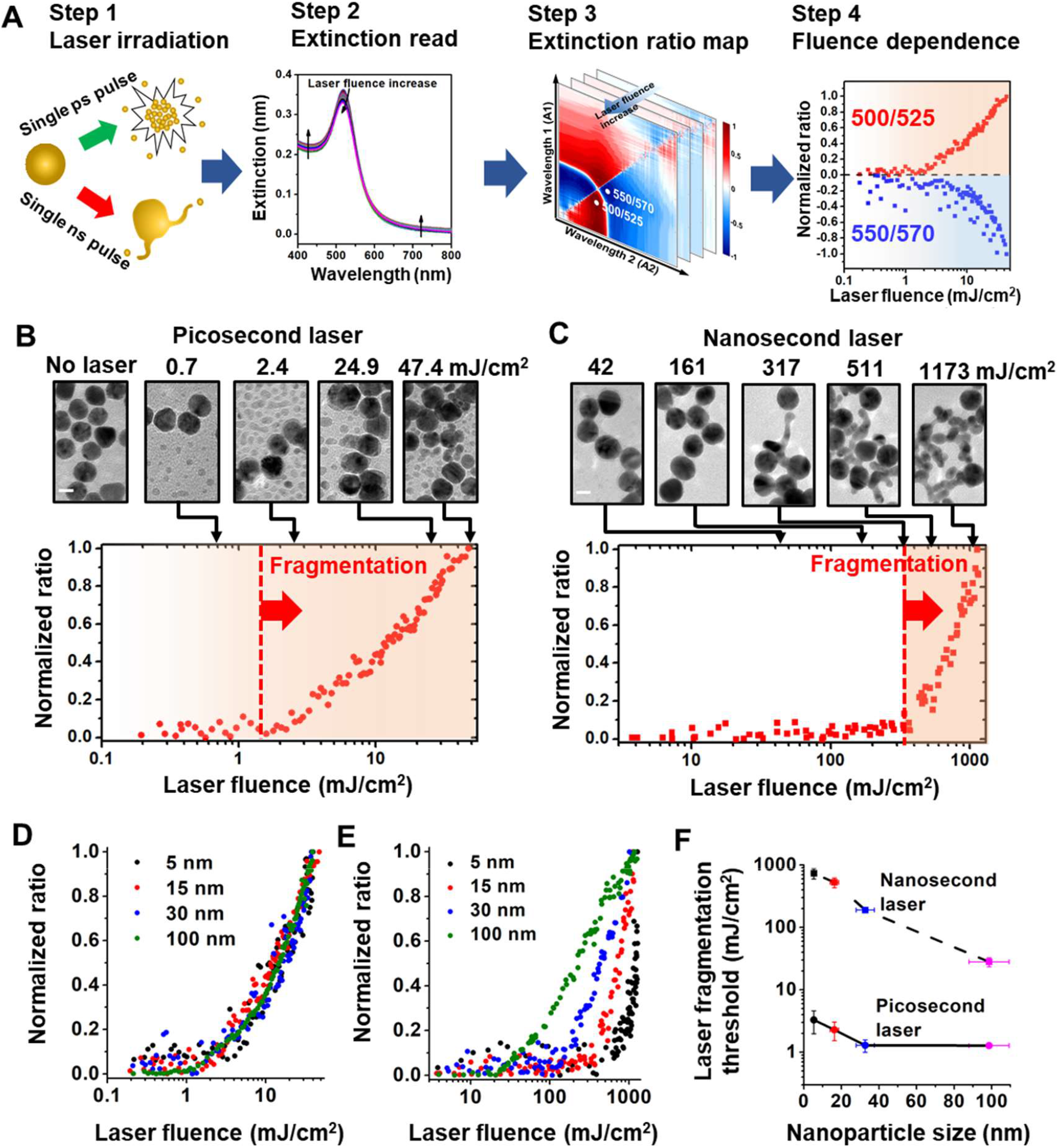
Fragmentation threshold fluences for gold nanosphere (AuNP) by laser irradiation. (A) Experimental and analysis procedure for fragmentation of AuNPs. (B-C) Extinction analysis and TEM images for fragmentation of AuNPs (diameter = 15 nm) by picosecond laser (B, Full width half maximum, FWHM = 28 ps) and nanosecond laser (C, FWHM = 6 ns). For picosecond laser, the normalized ratio at 500/525 nm shows increases when fluence is large than 1 mJ/cm^2^. For nanosecond laser, the ratio (450/500 nm) increase occurs when laser fluence is higher than 300 mJ/cm^2^ as the fragmentation threshold. TEM images support extinction analysis for fragmentation. (D-E) Extinction analysis for different particle sizes under picosecond laser (D) and nanosecond laser (E). (F) Threshold fluences of fragmentation under picosecond and nanosecond laser for different particle sizes.

Next, we examined nanoparticle fragmentation under ps and ns laser for a large range of AuNP particle sizes (5 nm to 100 nm). We compared the fluence dependence for extinction ratios with different particle sizes (Figure 1D-E and S2). We found that for ps laser-induced fragmentation the laser fluence threshold occurred in the range of 1-2 mJ/cm^2^ for all particle sizes (Figure 1D and 1F), indicating that the threshold for ps laser induced-fragmentation was weakly dependent on particle size. In contrast, the threshold fluence of fragmentation for ns laser exhibits a clear dependence on the particle size (Figures 1E and 1F), a 30-fold decrease from 600 mJ/cm^2^ for 5 nm AuNPs to 20 mJ/cm^2^ for 100 nm AuNPs. Finally, the thresholds for fragmentation observed for ps laser irradiation (1-2 mJ/cm^2^) were approximately 1-2 orders of magnitude lower than those observed for ns laser irradiation (20-600 mJ/cm^2^, Figure 1F).

### Picosecond laser excitation generates a bimodal nanoparticle size distribution that is reflected in the spectral analysis

To investigate the morphology of fragmentated nanoparticles under ps laser pulse, TEM images were analyzed to obtain the histograms for the size distribution of the AuNPs with different nominal sizes (5 nm, 15 nm, and 100 nm for Figure 2A-C) before and after exposure to single ps laser pulse. After laser irradiation, small fragments were detached from original particles. These newly generated daughter particles are mostly spherical but much smaller than parent particles.

**Figure 2.**
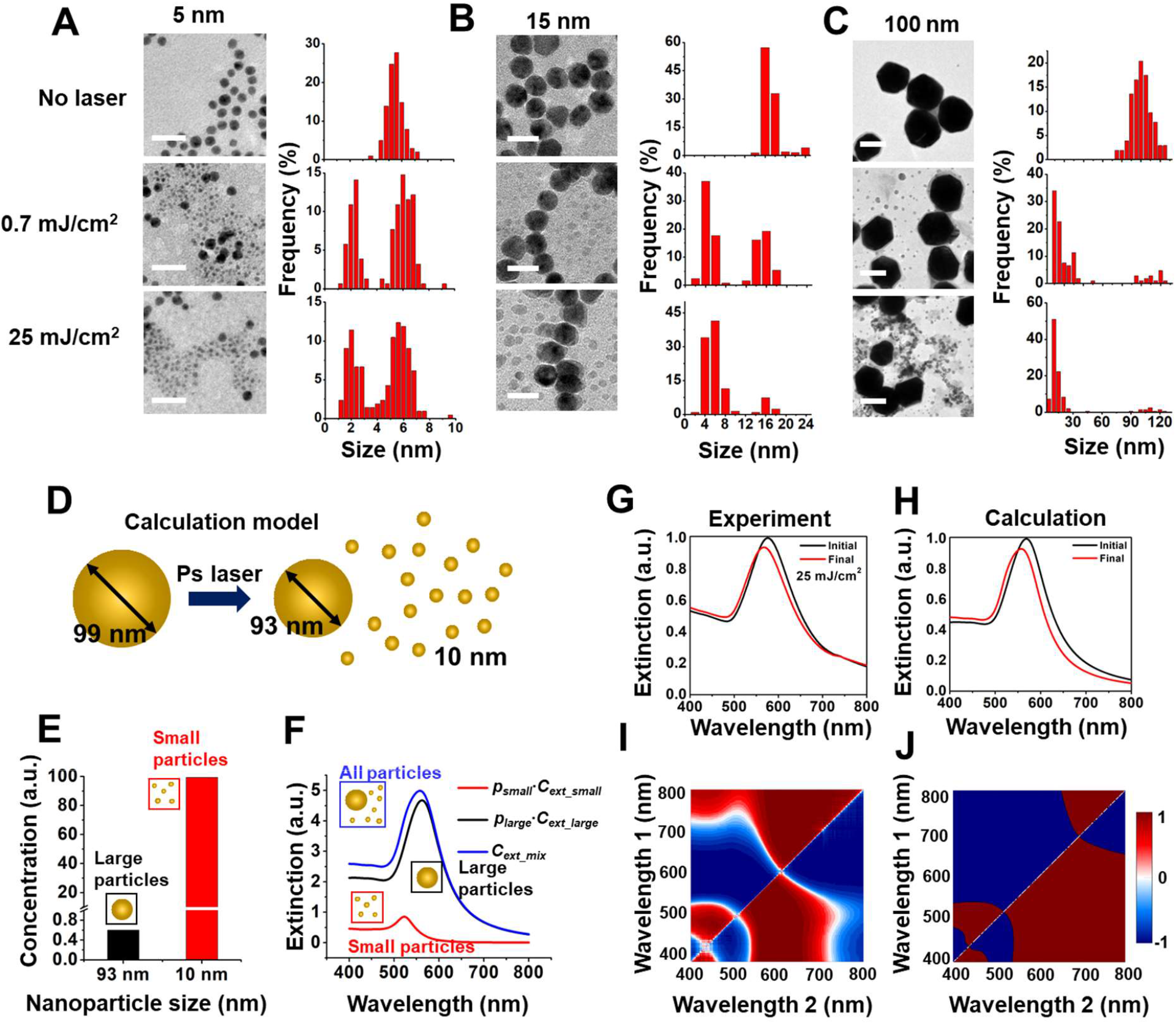
Bimodal size distribution for fragmentation under picosecond laser. TEM images and particle size distribution before and after picosecond laser irradiation for (A) 5 nm, (B) 15 nm and (C) 100 nm AuNPs. Scale bars are 20 nm for TEM images in (A) and (B) and 100 nm for TEM images in (C). Small particles with a relatively uniform size form after picosecond laser pulse irradiation. Higher laser energy (0.7 mJ/cm^2^ to 25 mJ/cm^2^) increases the number of small particle formation. (D) Schematic for modelling extinction spectrum change induced by AuNP size reduction. The initial size of particle is 99 nm and fragment sizes are 93 nm and 10 nm. (E) Concentration of small particles (10 nm) is much higher than that of large particles (93 nm) after laser irradiation. (F) Contributions of small particles (*p*_*small*_·*C*_*ext_small*_) and large particles(*p*_*small*_·*C*_*ext_small*_) to collective extinction (*C*_*ext_mix*_). (G-H) Experimental and predicted extinction spectra before and after laser fragmentation. (I-J) The extinction ratio maps from experiment and Mie theory calculation.

The population of daughter particles increases with laser fluence, and the size of daughter particles increases with the original particle size. For instance, after ps laser irradiation, the average diameters of daughter fragments are 2 nm, 5 nm, and 10 nm for AuNPs with original sizes of 5 nm, 15 nm, and 100 nm, respectively. We calculated the extinction spectra based on the size analysis of TEM images for 100 nm AuNPs, under the assumption that the change of total volume of AuNPs induced by fragmentation is negligible (Figure 2D). We calculated the concentration of daughter particles which have diameters of 93 nm and 10 nm (Figure 2E). Because the concentration of 10 nm AuNPs is much higher than that of 93 nm AuNPs (Figure 2E), the contribution of 10 nm AuNPs to the collective extinction is evident and causes a blue shift of the extinction peak. (Figure 2F-H). The calculated extinction spectra of AuNPs before and after laser treatment are comparable with experimental data (Figure 2G-J), and the 2D ratio map for extinction shows similar trend between experiment and calculation (Figure 2I-J). Therefore, we were able to recover the spectral changes resulting from the nanoparticle fragmentation under ps laser.

### Picosecond laser pulse fragments nanoparticles below nanoparticle melting point and Coulomb instability

Next, we compared ps and ns laser-induced fragmentation against known mechanisms of nanoparticle fragmentation, including Coulomb instability (CI) and phase explosion (PT). The temperature evolution of a AuNP with a diameter of 15 nm was calculated by a two-temperature heat transfer model (TTM) under both ps and ns laser irradiation, at their respective laser fluence thresholds for fragmentation of 2.3 mJ/cm^2^ (ps laser) and 527 mJ/cm^2^ (ns laser) (Figure 3A&S5). Because the ns laser has a duration (6 ns) that is much longer than electron-phonon coupling time (few ps), electrons are in equilibrium with the phonons so that the electron temperature (T_*e*_) calculation can be neglected ^23, 39^. Under ns laser irradiation, the model predicts that the gold lattice temperature exceeds the boiling point at its peak, suggesting that ns laser induced fragmentation is likely due to the PT mechanism. Although the electrons in gold are heated up to a very high temperature (∼3000 K) during ps laser irradiation, the gold lattice temperature is predicted to be well below the melting point, eliminating the possibility for the PT mechanism. To investigate whether CI contributes to nanoparticle fragmentation, the Rayleigh stability factor (X) was used to estimate the balance between the Coulomb force and the surface tension (Figure S5). The particle is unstable due to CI when the X factor is larger than 0.3 ^40^. For instance, the X factor of AuNPs with a diameter of 45 nm exceeds 0.3 when the fluence of the ps laser is between 17 and 18 mJ/cm^2^ (Figure S5C), indicating the fluence threshold for CI. For AuNP with a diameter of 15 nm, to reach the CI threshold, the fluence of the ps laser would need to be 10 times larger than the fragmentation threshold. Therefore, the ps induced particle fragmentation cannot be explained by the CI mechanism.

**Figure 3.**
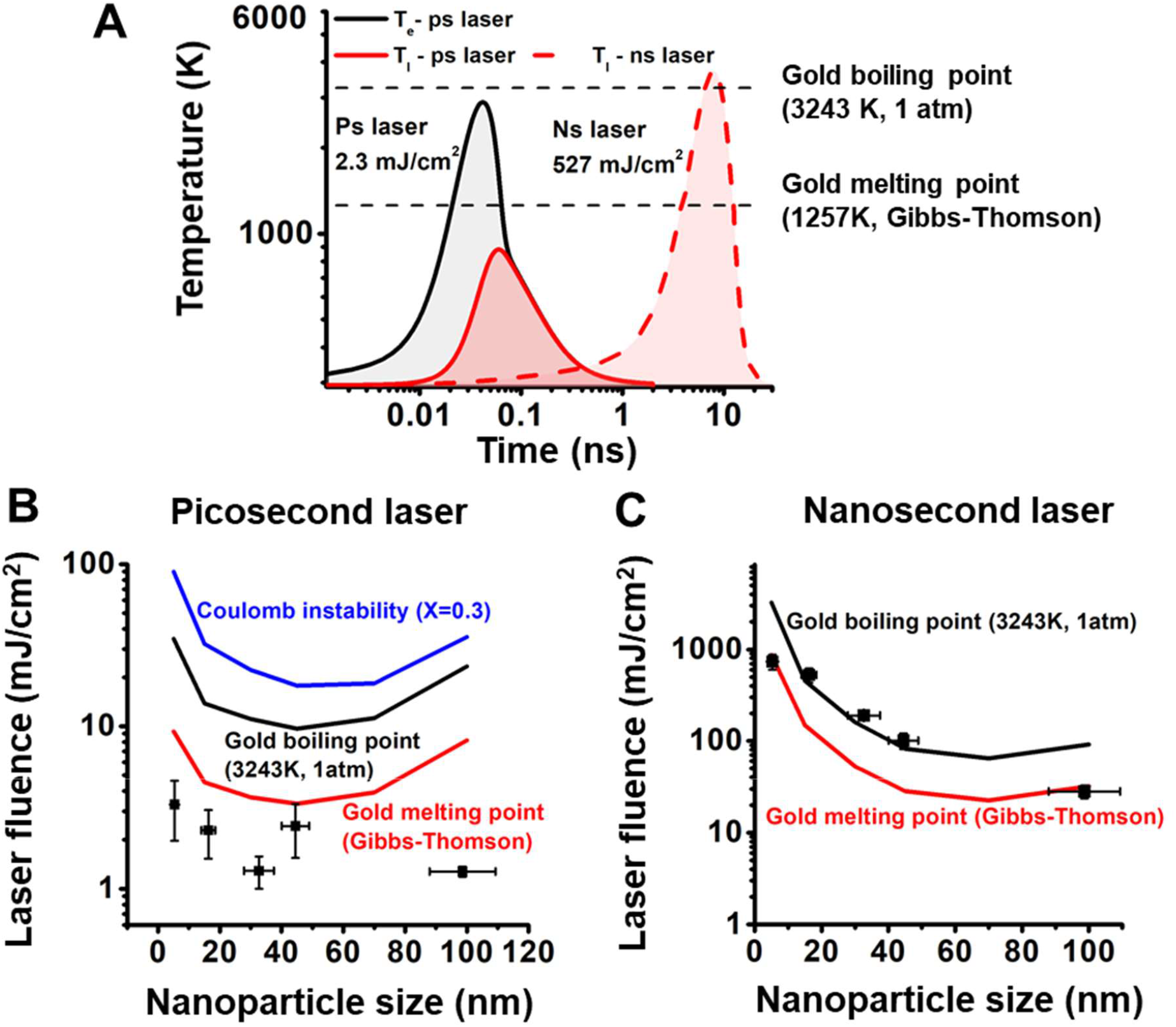
Mechanism of laser-induced fragmentation under picosecond (ps) and nanosecond (ns) laser. (A) Temperature evolution for 15 nm AuNP under ps and ns laser at fragmentation threshold fluences. Gold melting point and boiling point are marked with dashed line in the plot. (B-C) Comparison of the particle fragmentation threshold with gold melting and boiling point, and Coulomb instability for ps laser (B) and ns laser (C). The fluence of Coulomb instability achieves when Rayleigh instability factor (X) reaches to 0.3.

We further examined the fragmentation across a range of nanoparticle sizes. Varying nanoparticle size affects the plasmonic heating process in two ways. First, the absorption cross section is strongly dependent on particle size, causing different levels of heat generation for different particle sizes. Second, the heat dissipation from AuNP to water is dependent on the surface-to-volume ratio of the particle ^10^. Small particles have larger surface-to-volume ratio which makes the heat transfer more efficient than for larger particles. The temperature of a AuNP reflects the balance between these two factors. For instance, the 45 nm AuNPs achieve the highest particle temperature under ps laser heating, while the 70 nm AuNPs have the highest NP temperature under ns laser heating (Figure S6). This demonstrates that the duration of laser irradiation can have a significant effect on the nanoparticle heat transfer, allowing different thermophysical processes to occur for different sized particles.

Next, we systematically examined the threshold fluences for different particle sizes (Figure 3B&C). Interestingly, the threshold fluence for fragmentation under the ps laser is below the melting point for all particle sizes (5∼100 nm). Since both PT and CI explains fragmentation of AuNPs above the melting point, these two mechanisms cannot be responsible for the ps laser-induced fragmentation. On the other hand, the threshold fluence for fragmentation under ns laser shows a similar trend as the gold boiling point ^41^. Therefore, we conclude that the PT mechanism is the dominant mechanism for fragmentation of AuNPs under ns laser heating. We also tested the effect of sodium dodecyl sulfate (SDS) on the particle fragmentation. We compared the fragmentation behavior of 15 nm AuNPs in pure water with that in the SDS solution at a concentration of 90 mM, which is higher than its critical micelle concentration (8.9 mM) (Figure S7). Previous study demonstrated that the negative charge of SDS micelles is able to stabilize holes formed upon ionization, resulting in an increase in the ionization yield ^42^. If the fragmentation is induced by thermionic emission under the CI mechanism, the fragmentation should be enhanced by the addition of SDS. However, our results shows that the addition of SDS hindered the fragmentation under ps laser, suggesting that the fragmentation under ps laser does not originate from laser ionization.

Lastly, we studied the contribution of thermomechanical force to the ps laser fragmentation using molecular dynamics (MD) simulation^43^. AuNPs with diameter of 5 nm and 15 nm can be fragmented when the laser intensity was higher than 40 and 20 mJ/cm^2^, respectively (Figure S8A&B). The fragmentation cases show the maximum gold temperature above the boiling point (Figure S8C&D). The nanoparticle heating leads to thermal expansion and a high mechanical stress within the nanoparticle (Figure S8E-G). The thermomechanical force causes movement of atoms and leads to a disordered lattice structure. In order to release the thermomechanical force, some unstable atoms are ejected from the particle, behaving as the fragmentation. The MD simulation demonstrated the thermomechanical force plays an important role in the phase explosion induced fragmentation. However, it is worth noting that our MD study demonstrates that particle fragmentation occurs only above the boiling point which does not agree with our experimental observations. Therefore, the thermomechanical stress was not sufficient to explain the anomalous fragmentation observed in ps laser experiments.

### Comparison with previous works

We summarized previous reported experimental conditions for ps laser induced plasmonic nanoparticle fragmentation in the Table S2. Most studies reported plasmonic nanoparticles can be fragmented by lasers with fluences in the range of 10∼30 mJ/cm^2^, which brought the particle temperature above the boiling point and suggested a phase explosion mechanism. The ps induced AuNP fragmentation at solid state in this study has threshold fluences lower than most reported data, which indicates an unknown mechanism.

### The effect of interfacial heat transfer

The heat transfer process at the interface of nanostructures is still an emerging area with many unknowns. The interface thermal resistance used in this study is obtained from previous measurement^44^. However, the interface heat transfer can be a function of temperature, interface curvature, solvent type, synthesis method, and nanoparticle morphology^45, 46, 47^. The variation of interface heat transfer can significantly affect the gold temperature, especially for ns laser^48^. Also, the phase change in water was not considered in the current model. Water can be heated and evaporated on the surface of the nanoparticle upon laser irradiation, generating what is known as a plasmonic vapor bubble. The vapor bubble formation will modulate the resonance condition of a AuNP and alter the heat transfer. The nanobubble forms when the water temperature reaches the spinodal temperature (about 85% of the critical point), which is within the range of our study^49^. These factors need to be considered in the future to achieve a more accurate prediction.

In our experiment, we utilized citrate coated gold nanoparticle to conduct the laser fragmentation experiment. Some studies demonstrated that the citrate molecules can play an important role in the particle shape evolution during laser irradiation^50^. Using bare AuNPs to study the laser induced fragmentation can exclude the effect of surface ligands^33^. So far, it is still not clear that how the surface ligands affect the laser-nanomaterial interaction as well as the heat transfer process. It is possible that the citrate coating alters the interfacial thermal resistance and cause a temperature difference in gold than the model prediction^48^.

### Possible contributions from near-field ablation and surface melting

Near-field ablation is another mechanism responsible for nanostructure fragmentation. For example, Plech *et al*. demonstrated that AuNPs can be fragmented by near-field ablation below the melting point under fs laser irradiation^28^. Near-field ablation is highly related to the polarization of the incident light and the morphology of nanoparticles^29^. For example, Yan *et al*. studied the fs laser-induced near-field ablation of Ag nanorods and found that the ablation is highly related to the electric-field distribution. Although it was reported that sub-ps laser can also induce near-field ablation^30^, most applications of near-field ablation require femtosecond laser. Another important factor that contributes to the change of particle morphology is particle surface melting^51^. Importantly, surface melting of gold nanostructures can be observed at a much lower temperature than the melting point of the bulk gold. For instance, Inasawa *et al*. demonstrated that AuNPs can be reshaped and fragmented under 355 nm ps laser irradiation^51^. Under those conditions the laser energy was insufficient to melt the whole particle, suggesting that the observed change in morphology was induced by surface melting. Petrova *et al*. observed obvious shape transitions of gold nanorods under oven heating of less than an hour at 423K (∼32% of the melting point of bulk gold). Similarly, Plech *et al*. demonstrated the surface melting of AuNPs supported on a surface occurs at 377 K (30% of the melting point of bulk gold)^52^. The temperature of AuNP (30 nm) at the ps laser fragmentation threshold fluence is around 700 K, which is well above the surface melting point. As such, we expect that during ps laser heating, the surface of AuNPs becomes molten while the core of particle remains still solid. With this in mind, one hypothesis to explain our observation that ps laser-induced fragmentation occurs below the (whole) particle melting point is that the combination of surface melting and near-field enhancement induces small particle ejection from the surface of the AuNPs. However, further studies are required to fully uncover the mechanism of fragmentation under ps laser irradiation.

## CONCLUSIONS

In this study, we investigated the fragmentation of AuNPs under ps and ns laser with experiments and numerical simulations. The behaviors of fragmentation induced by ps and ns laser show obvious differences in aspects of the threshold fluence and morphology change. Whereas the ns laser induced fragmentation can be explained by the PT mechanism, neither PT nor CI mechanisms can fully explain the ps laser induced fragmentation which occurs while the AuNPs are still in the solid state (Figure 4). Although the molecular dynamic simulations suggested that the mechanical stress may play a significant role in ps laser-induced fragmentation under high laser intensity, it could not explain ps laser-induced fragmentation below the AuNP melting point. This study reveals a previously unknown regime of nanoparticle fragmentation.

**Figure 4.**
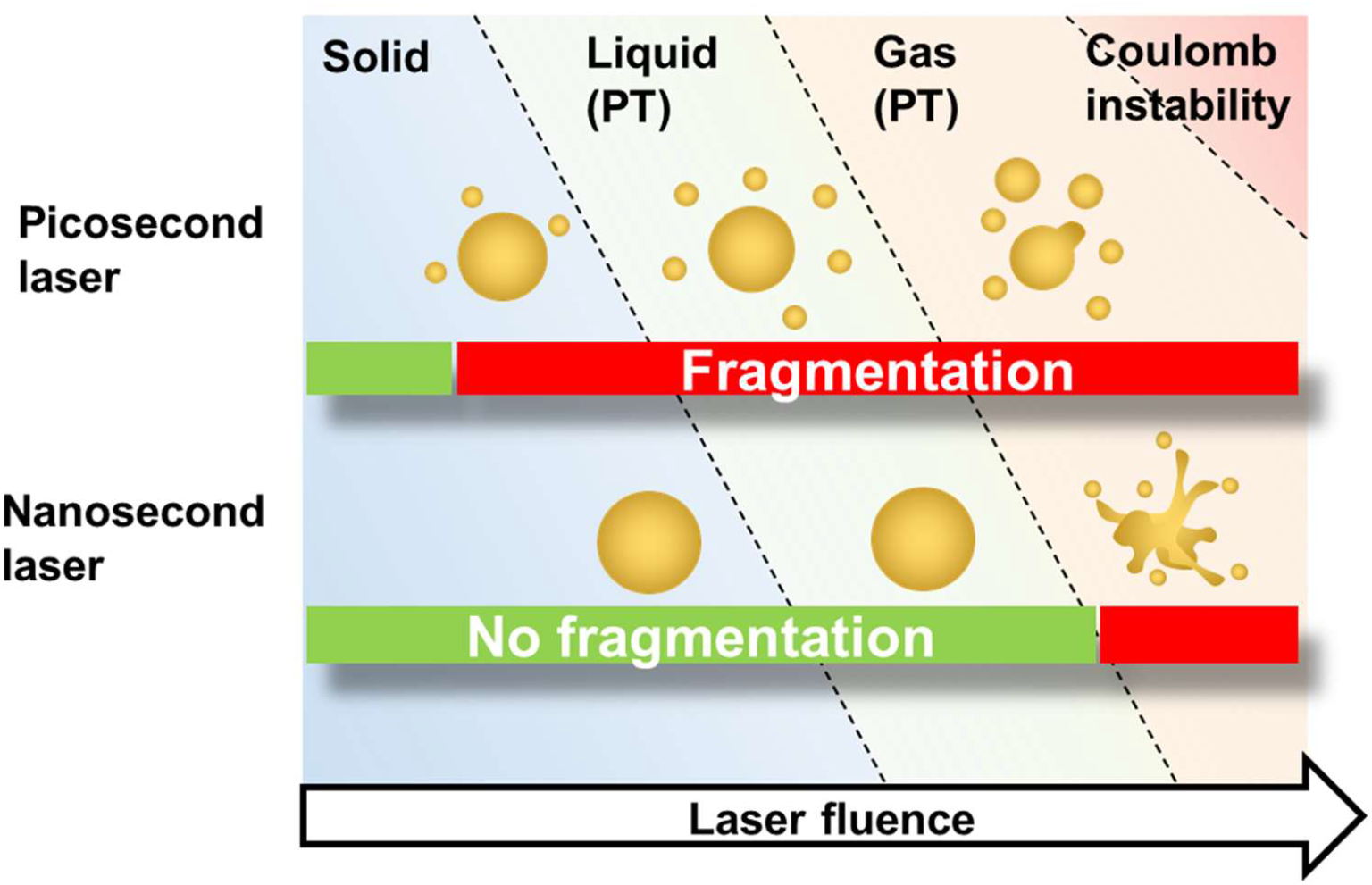
Proposed schemes for laser-induced fragmentation under picosecond and nanosecond laser pulses.

## Materials and Methods

### Nanoparticle synthesis

All chemicals were purchased from Sigma-Aldrich unless otherwise indicated. 15 nm AuNPs was synthesized with the Frens’ method following the previous report^53^. All glassware was soaked in aqua regia for 30 mins and rinsed with the Millipore water before using. 98 mL Millipore water and 1 mL HAuCl_4_ (25 mM) were added to 250 mL Erlenmeyer flask and brought to a boil, following the rapid addition of 1 mL trisodium citrate (112.2 mM). The solution was heated for 10 minutes and cooled down to the room temperature. Particles were concentrated with centrifugation (10 krcf, 30 minutes), and stored in 4°C environment until experiment.

30, 45, 100 nm AuNPs were synthesized with the seed-growth method with 15 nm AuNPs as seeds^54-55^. Briefly, 15 nm AuNP (2.23 nM), HAuCl_4_ (25 mM) and trisodium citrate (15 mM) solution was mixed and brought to the volume of 100 mL with Millipore water. Under rigorous mixing, hydroquinone (25 mM) solution was added into the vortex swiftly. The solution was stirred for half an hour under the high-speed stirring and then changed to low-speed stirring overnight. The volume of each solution is dependent on NP sizes and can be found in previously published paper^55^. Particles will be concentrated by centrifugation and stored in 4°C environment until experiment. The centrifugation speed and time were dependent on particle sizes.

5 nm AuNPs was synthesized following a previously reported procedure^56^. The reduction solution was prepared by mixing 16 mL water, 4 mL trisodium citrate (1% w/v), 1 mL tannic acid (1% w/v) and 1 mL potassium carbonate (3.26 mg/mL). The reaction is started by injecting reduction solution to 80 mL HAuCl_4_ solution (312.5 µM). The reaction was kept under 60 °C for half hour and then brought to 90 °C for 10 minutes.

All nanoparticles are cooled to room temperature. Sizes of AuNPs are characterized with transmission electron microscope (TEM) and dynamic light scattering (DLS, Malvern Nano ZS). To characterize the AuNP sample with TEM, AuNP solution (10 µL) was dropped on TEM grid and air dried. The TEM imaging was taken in JOEL-2100 (100 keV). Imaging analysis were conducted in ImageJ software.

### Laser beam size measurements

The ns pulsed laser is generated from a Quantel Q-smart 450 Nd:YGA laser. The ps pulsed laser is generated from EKSPLA PL 2230 Nd:YGA laser. The laser beam sizes are calibrated with blade-edge method following a previous report^57^. We firstly fixed a razor blade on a translation stage in front of the laser beam. Next, the laser energy was monitored while a blade edge is gradually blocking the laser beam. The curve shows the relation between the blade position and laser power was used to calibrate the gaussian beam size using the relation below.

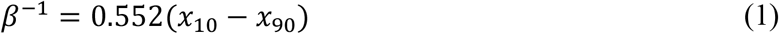

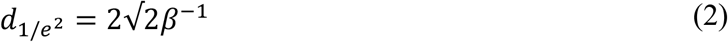

where *β*^-1^ is radius where laser intensity drops to 1/*e* of the maximum intensity in the center of the beam. And 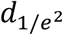 is the diameter where intensity drops to 1/*e*^2^ of the maximum intensity.

### Laser-induced nanoparticle fragmentation

Before the laser-induced fragmentation experiment, all AuNP are diluted to the OD=0.3 at 532 nm. Then AuNP solution is added to a 96 well-plate (115 µL/well). We irradiated samples well by well with increasing laser fluence. The pulse energy was monitored simultaneously with a beam splitter which can reflect portion of the beam. The transmission and reflection were measured by an energy meter, and the linear relation between transmission and reflection was calculated in Excel. After exposing to single laser pulse, samples (110 µL/well) were transferred to 384 well plate. We used a plate reader (BioTek Synergy 2) to measure the extinction spectrum from 400 nm to 800 nm.

### Data processing and extinction analysis

The extinction spectrum for different laser fluences was organized in Excel. The data was then processed with MATLAB 2019a. The extinction spectrum is a function for both wavelength (*λ*, from 400-800 nm) and laser fluence (*F*), as shown in Eq. 3. Here, *A* is extinction spectrum.

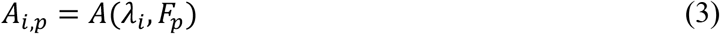

The extinction spectrum of AuNP after laser was treated as a vector [*A*]. The extinction ratio, as a matrix [*R*], was calculated by Eq. 1. For each laser frequency, we can obtain a matrix [R].

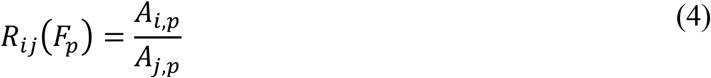

We then normalized the extinction ratio by scaling between 0 to ±1 with Eq. 4 where F = 0 indicates the control sample without laser irradiation, and *F* = *F*_*q*_, refers to the sample with the largest ratio change among all laser fluences.

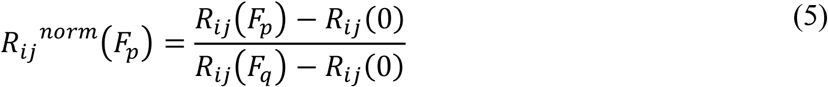

The threshold fluence was defined as where the normalized extinction ratio starts to increase. We used a piecewise fitting model to fit the data and found the optimized breaking point which indicates the threshold fluence^58^.

### Mie theory calculation

The extinction of AuNP before and after fragmentation under ps laser was calculated by Mie theory with MNPBEM Matlab toolbox^59^. We assume the gold density is constant for particles with different sizes and there are only two discrete sizes of particles according to TEM images. The average diameters (d) of nanoparticles were obtained from TEM imaging analysis. The collective extinction spectrum of AuNP sample (*C*_*ext_mix*_) can be calculated by eq. 6-8.

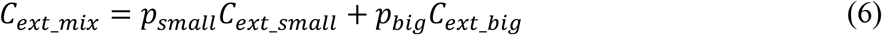

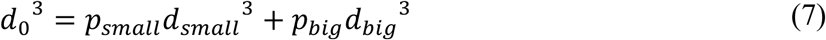

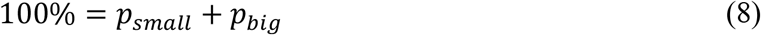

where *C*_*ext*_ is the extinction cross section, *p* is portion of particles with different sizes. Subscripts “0”, “small” and “big” indicates the particle before laser, small fragments, and big remained particles, respectively.

### Two-temperature heat transfer model

In this paper, the shortest pulse we used was ps laser which should be sufficiently long to neglect electron relaxation process (in several femtoseconds). Therefore, the parabolic two-step heat conduction model (P2T) is sufficient for our estimation of the temperature evolution^39^. TTM describes that the electron subsystem in the AuNP obtains energy from laser, and then transfer energy to lattice.

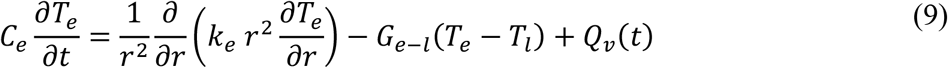

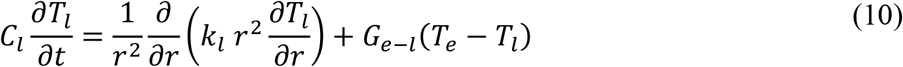

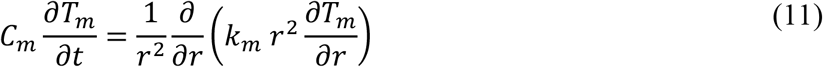

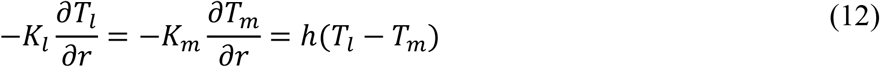

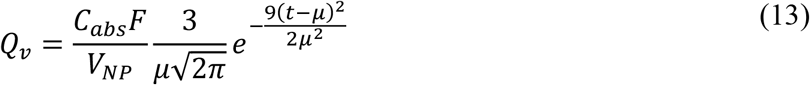

where the *Q*_*v*_ is volumetric heat source, *C*_*abs*_ is absorption cross section computed from Mie theory, *F* is laser fluence in mJ/cm^2^. The laser is treated as a gaussian pulse in the simulation. The total pulse duration was set as 6*σ* of the gaussian function. And the peak of the pulse occurs at *μ*. For ps laser (FWHM=28 ps), the *μ* is 35.67 ps. For ns laser (FWHM = 6 ns), the *μ* is 7.64 ns. Subscript letters “e”, “l” and “m” refer to electron, lattice and medium. *K* is thermal conductivity, *C* is specific heat and *G* is thermal coupling factor. The heat trasfer between water and nanoparticle has a finite interfacial thermal conductance, *h*. The value and expression of thermal properties are listed in table S1.

For ns laser heating, due to the relatively long laser duation, we neglected the electron-phonon interaction and only consider heat transfer by phonons. Therefore, tradiational one temperature model (OTM, heat conduction equation) and TTM leads to the same result.

The size dependent melting point for gold nanosphere is calculated through the Gibbs-Thomson relation:

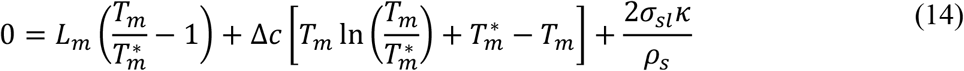

where *L*_*m*_ is the latent heat of gold during melting, *T*_*m*_ is the size dependent melting point, *T*_*m*_* is the melting point of the bulk gold, *σ*_*sl*_ is the interface tension between the solid and liquid gold, *κ* is the mean curvature, *ρ*_s_ is the solid gold density (19300 kg m^-3^). Δ*c* is the difference between the specific heat of solid and liquid gold (34 J kg^-1^ K^-1^)^60^.

### Molecular dynamics simulation

The process of gold nanoparticle fragmentation induced by ps laser irradiation was numerically investigated using the LAMMPS molecular dynamics package^61-62^. Single AuNPs with diameters of 5 and 15 nm were immersed in a cubic water box with side dimension of 10 and 20 nm, respectively. The system was initially equilibrated to ambient temperature (300K) and pressure (1atm) under NPT ensemble. Laser heating was then simulated by applying a uniform heating power to the AuNP atoms in order to rapidly heat the AuNP, followed by a final time period during which the system was allowed to return back to ambient conditions. The periodic boundary condition (PBC) was applied along every direction for all the simulations. For solvation, the TIP3P water model was adopted with the interaction energy computed as follows^63^,

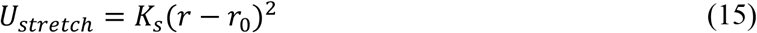

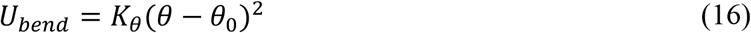

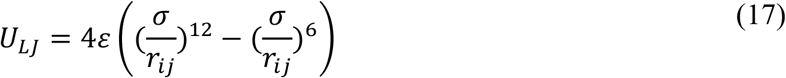

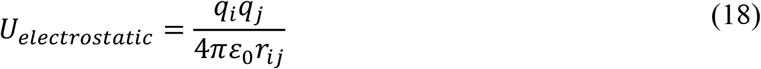

The *U*_*stret*_ and *U*_*bend*_ compute the bonded interaction for bond stretching between neighbor atoms and angle bending between neighbor bonds, respectively. The nonbonded interaction composed of two terms. First, the van der Waals interaction, *U*_*LJ*_, which was described by 12-6 Lennard-Jones (LJ) potential. Second, the *U*_*electrostatic*_ accounts for the Columbic interaction between charged atoms. The parameters are listed in Table S1.

For the interaction among gold atoms, the embedded atom method (EAM) was adopted using the parameters developed by Grochola *et al*.^64^. The total energy of each atom was computed as follows,

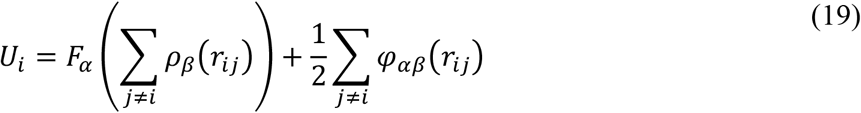

where *F*_a_ is the embedding energy which is a function of the atomic electron density *ρ*_β_, *φ*_αβ_ is the pair potential interaction with *α* and *β* denoting each pair of atom type. Since we only have one metal atom, Au, in the system, *α* and *β* would be identical here.

### Models for the thermionic emission and the Coulomb instability

When electrons exceed the critical energy state, they can be ejected from gold atoms and leave the atom with positive charges, known as electron ejection. The multiphoton stimulation and photoelectronic effect can be neglect for ps and ns laser heating ^23^. Therefore, the thermionic emission is the only mechanism for electron ejection in our model. The number of ejected electrons from single particle was calculated by equation:

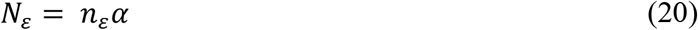

where *n*_*E*_ is electrons that ejected per atom and α is the total number of single AuNP. *n*_*E*_ can be calculated by following equation:

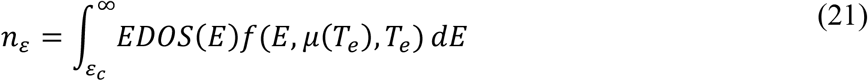

where EDOS is electron density of states function, *ε*_*c*_ is critical energy state (10.23 eV). *f* is Fermi-Dirac distribution:

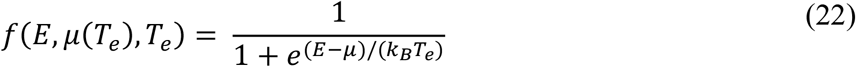

where *E* is energy level (in eV), *μ*(*T*_*e*_) is chemical potential, and *T*_*e*_ is electron temperature. *α* can be calculated by following equation:

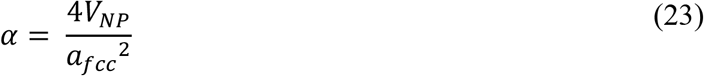

We assumed the AuNP is spherical and *V*_*NP*_ is volume of the AuNP. *a*_*fcc*_ is the basic length of the unit cell in fcc crystal, 4.08 Å for gold.

Lastly, we calculated Coulomb instability using Rayleigh instability and metal fission model. The Rayleigh instability factor *X* was calculated by following equation:

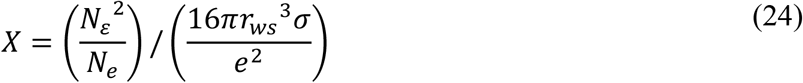

where *N*_*e*_ is total number of free electrons in gold. We assume every atom contributes one electron, thus, *N*_*e*_ = α. r_*ws*_ is Wigner-Seitz radius, *σ* is surface tension of gold, and e is elementary charge. The parameters used in this paper are listed in Table S1.

## ASSOCIATED CONTENT

### Supporting Information

Supporting information contains following content: Figure S1. Laser beam profile measurement. Figure S2. Extinction analysis for different particle sizes. Figure S3. Material properties for two temperature model. Figure S4. Temperature evolution for 15 nm AuNP under the ns laser in figure 3A. Figure S5. Parameters for the Rayleigh instability model. Figure S6. Volumetric heating factor and the maximum temperature of gold lattice.

Figure S7. Influence of sodium dodecyl sulfate (SDS) on picosecond laser fragmentation. Figure S8. Molecular dynamics simulation for fragmentation of AuNP. Table S1. Parameters in TTM model and MD simulation. Table S2. Previous reported laser fluence of ps laser fragmentation for plasmonic nanoparticles. The following files are available free of charge. Supporting information (PDF)

## AUTHOR INFORMATION

### Corresponding Author

*

### Author Contributions

Conceptualization: Z.Q. & P.K. Experiments: P.K. Computational modeling: P.K., Y.W. & B.A.W. Writing-original draft: P.K. Writing-review and editing: B.A.W., Y.W., J.R., A.P., & Z.Q.

## ACKNOWLEDGMENT

P.K. thanks for the useful discussion with Mr. Napat Dawkrajai. Funding: Research reported in this publication was partially supported by National Institute of General Medical Sciences of the National Institutes of Health under award number R35GM133653, and by the National Science Foundation under Grant No. 1631910. The content is solely the responsibility of the authors and does not necessarily represent the official views of the funding agencies.

