## supporting information for "Nanoparticle fragmentation at solid state under single picosecond laser pulse stimulation"

**This PDF file includes:**

Figs. S1 to S7
Tables S1
References (1 to 11)

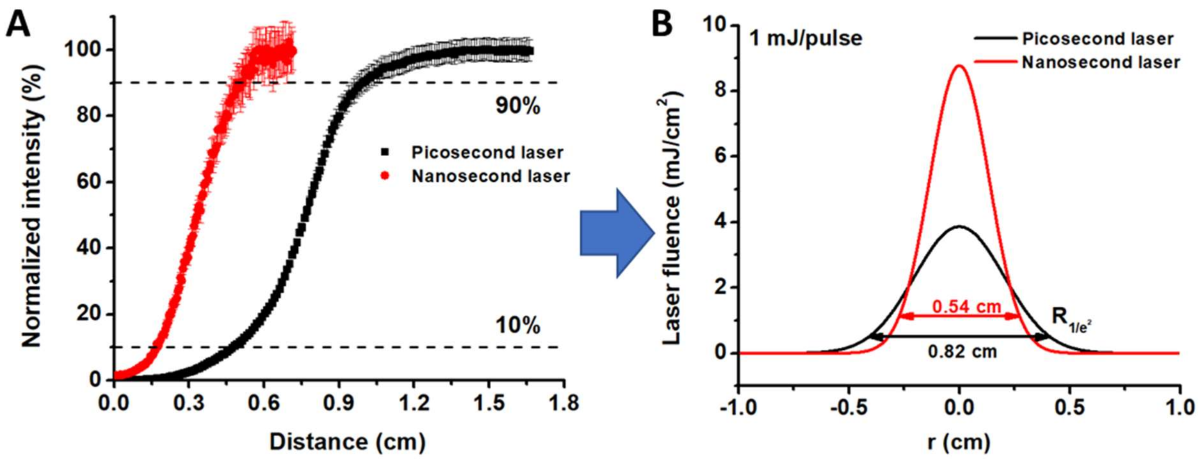


**Figure S1. Laser beam profile measurement** (A) Blade-edge measurement of laser beam profile for nanosecond and picosecond laser beam. (B) Gaussian laser beam profile for nanosecond and picosecond laser.

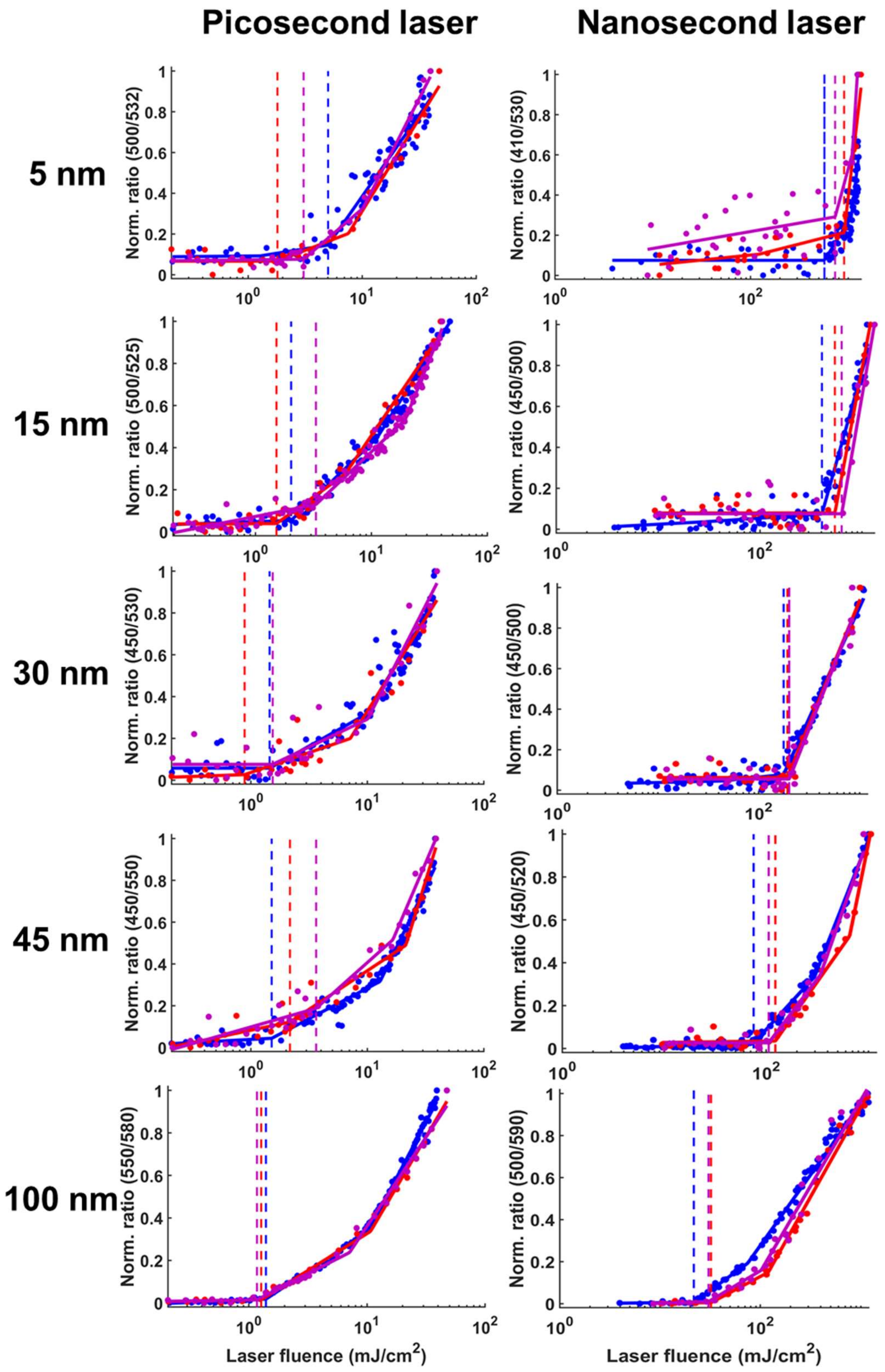


**Figure S2. Extinction analysis for different particle sizes.** The left panel is picosecond laser induced fragmentation. The right panel is nanosecond laser induced fragmentation. Different color of data is obtained from different experiments.

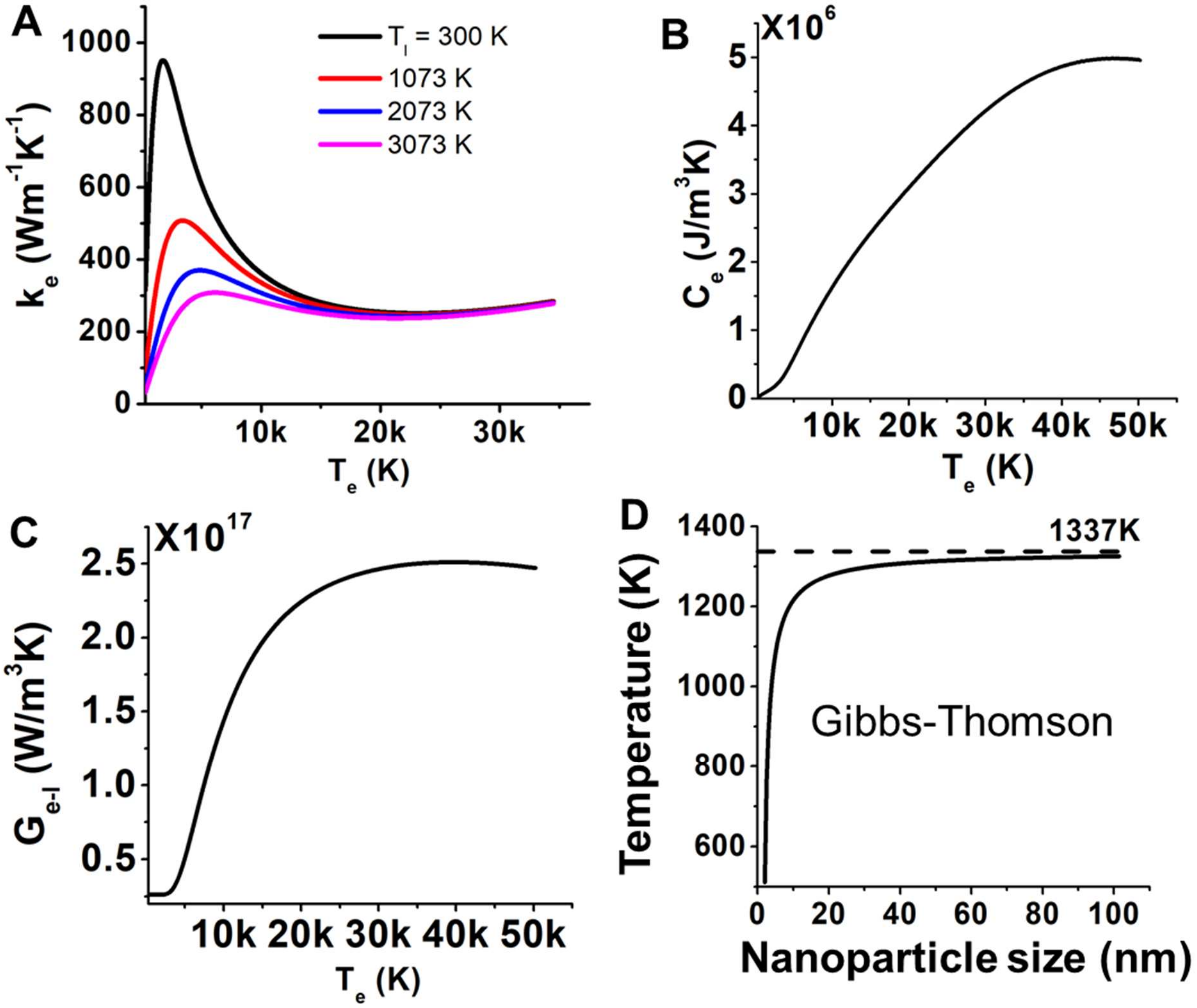


**Figure S3. Material properties for two temperature model.** (A) Thermal conductivity of electrons changes with electron temperature ($T_e$) and lattice temperature ($T_l$). (B) Specific heat of electron. (C) Electron-phonon coupling factor ($G_{e-l}$). (D) Size dependent melting point for gold nanoparticles calculated by Gibbs-Thomson equation.

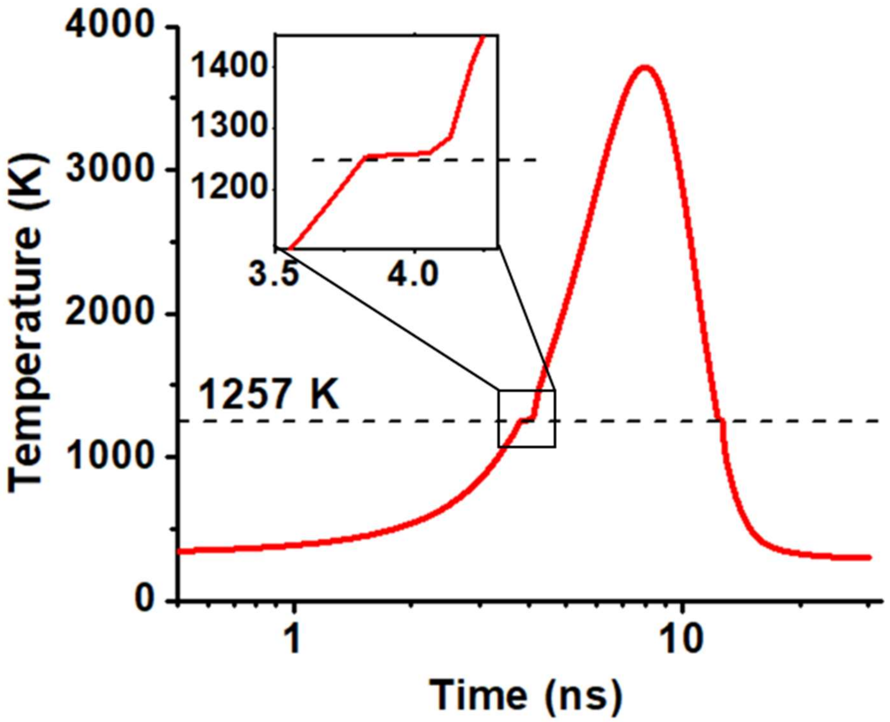


**Figure S4. Temperature evolution for 15 nm AuNP under the ns laser in figure 3A.** The phase transition occurs at 1257 K as the gold melts.

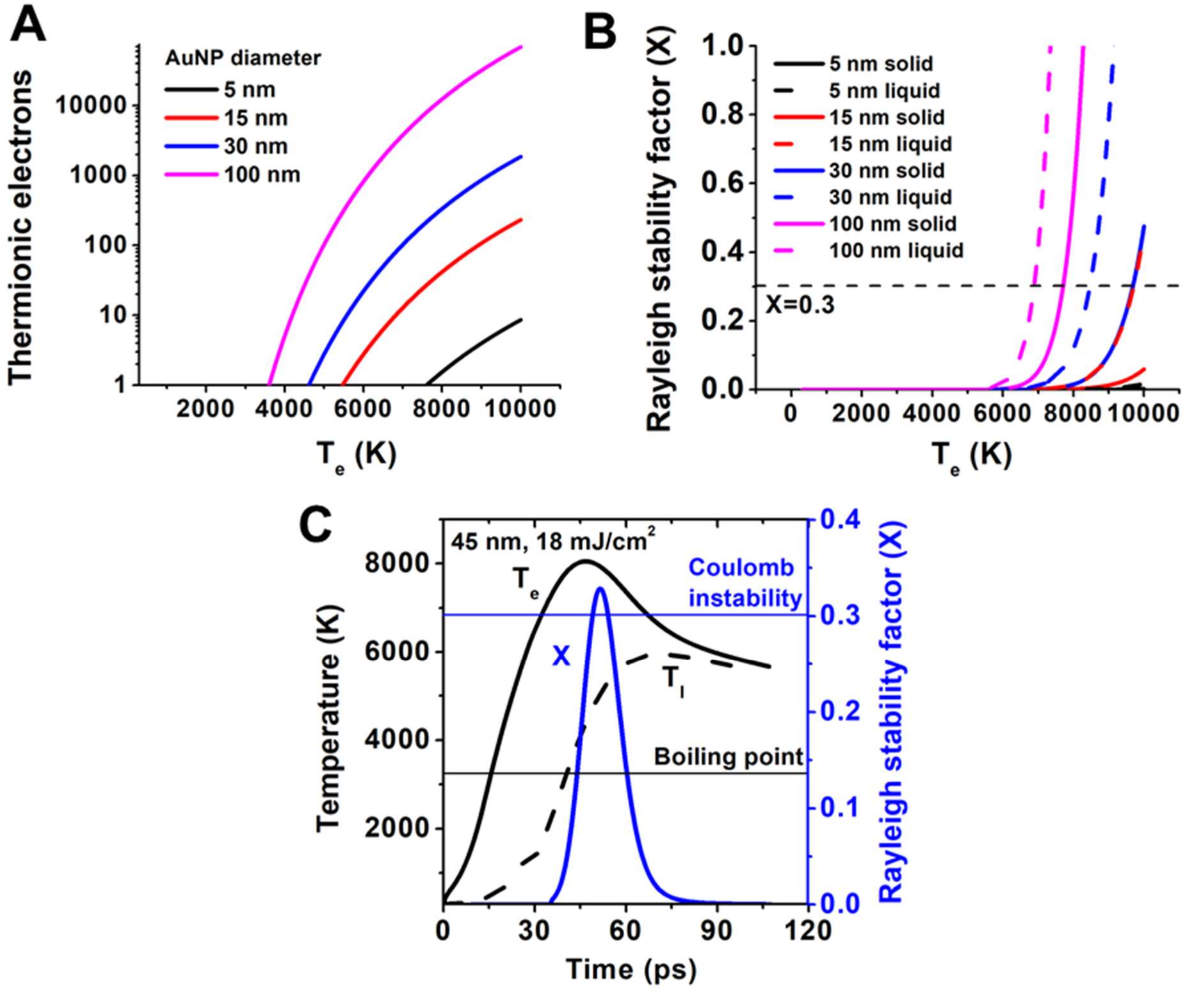


**Figure S5. Parameters for the Rayleigh instability model.** (A) The thermionic electrons and (B) Rayleigh stability factor (X) as a function of electron temperature ($T_e$). (C) Evolution of temperature and X for 45 nm gold nanosphere (AuNP). X is larger than 0.3 when laser fluence is 18 mJ/cm$^2$.

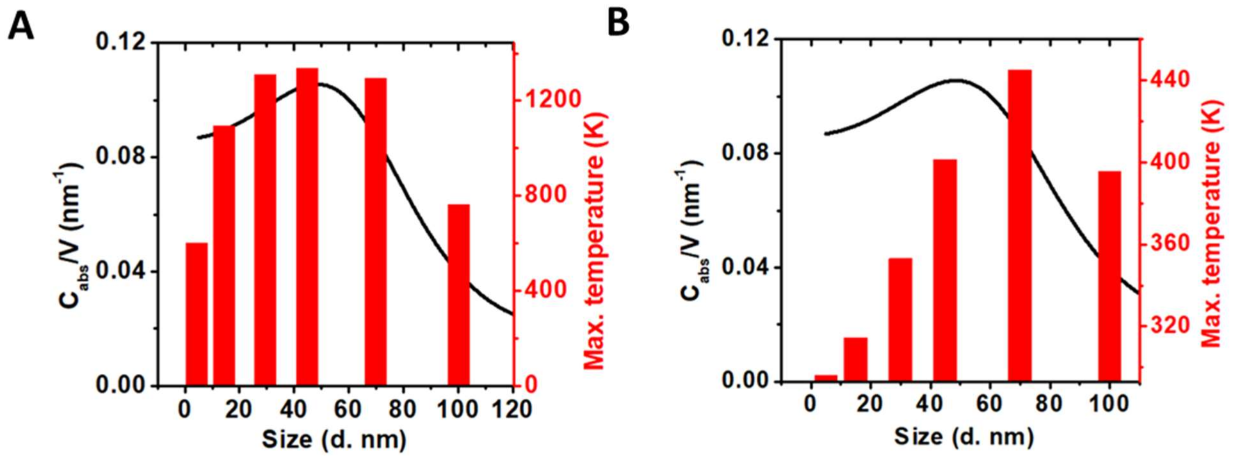


**Figure S6. Volumetric heating factor and the maximum temperature of gold lattice** for (A) picosecond laser and (B) nanosecond laser heating. $C_{abs}$ is absorption cross section area and V is volume of NP. $C_{abs}/V$ represents amount of light energy absorbed by the particle per volume. The laser fluence is the same for both cases (3.16 mJ/cm$^2$).

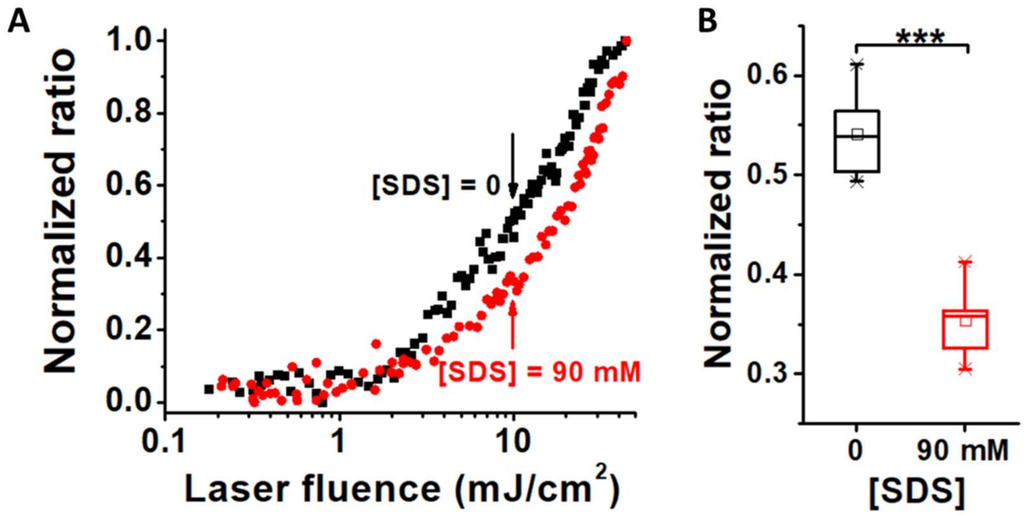


**Figure S7. Influence of sodium dodecyl sulfate (SDS) on picosecond laser fragmentation.** (A) Extinction analysis for picosecond laser ablation for 15 nm AuNP in 90 mM SDS solution (red dots) and in water (black dots). Picosecond laser ablation can be delayed by addition of SDS. (B) Comparison of normalized ratio at 10 mJ/cm$^2$ for AuNP ablation in water and in 90 mM SDS (n=6). - indicate maximum and minimum data, X indicates 99% and 1% confidence, □ represents the mean value, box and bars represent 25%, 75% confidence and the median value. *** indicates $p<0.05$.

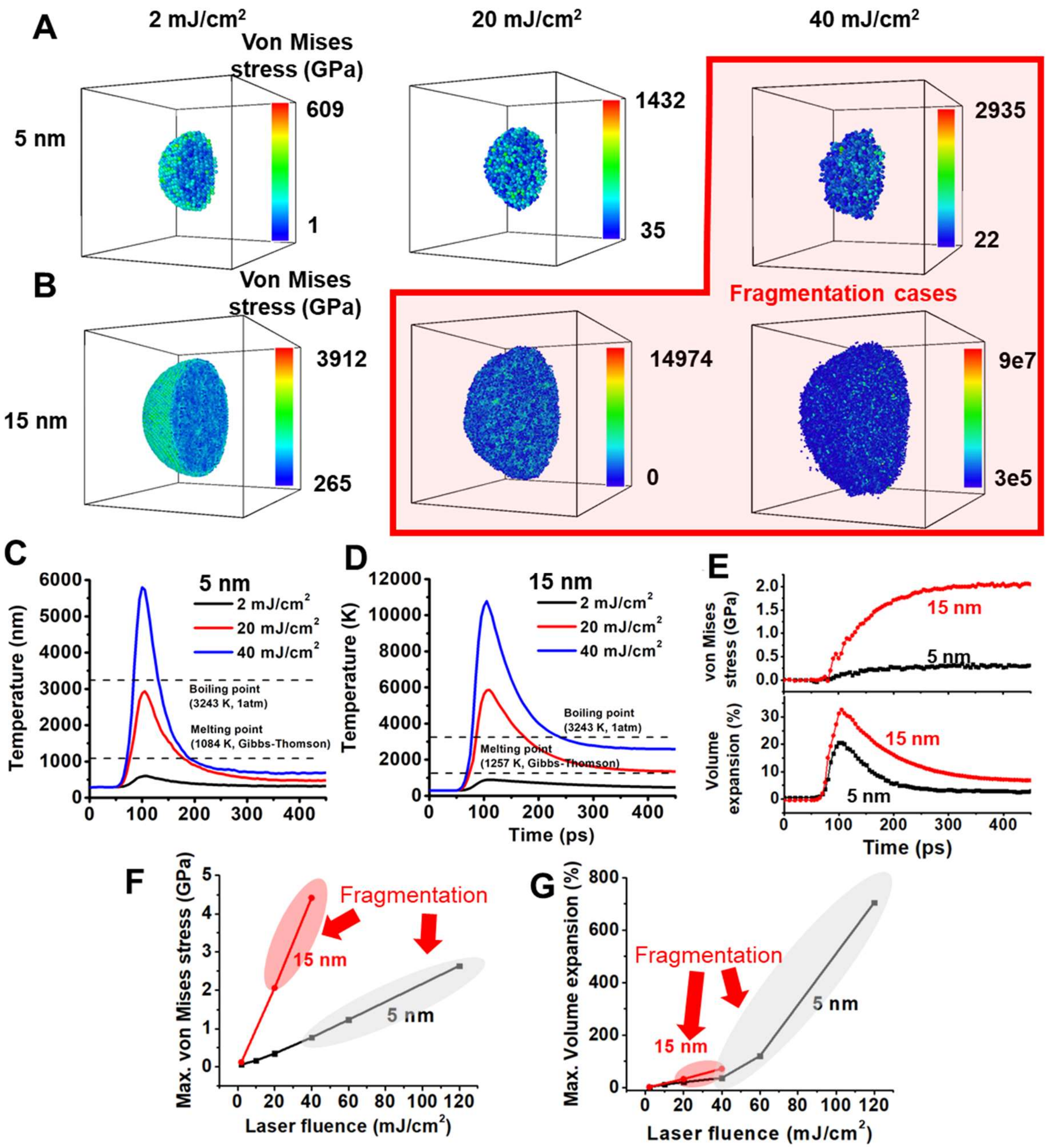


**Figure S8. Molecular dynamics simulation for fragmentation of AuNP.** The contour plot of von Mises stress of (A) 5 nm AuNP and (B) 15 nm AuNP with the cross section cutting through the center of AuNP when the particle reaches maximum temperature. Daughter particles were observed when laser intensity is larger than 40 mJ/cm$^2$ for 5 nm AuNP and 20 mJ/cm$^2$ for 15 nm AuNP. (C-D) The average gold temperature for (C) 5 nm AuNP and (D) 15 nm AuNP. The melting point and boiling point are marked with dashed lines. (E) The average von Mises stress and volume expansion for 5 nm (black) and 15 nm (red) AuNP with the laser intensity of 20 mJ/cm2. (F) Maximum von Mises stress for different laser intensities. (G) Maximum volume expansion for different laser intensities. The fragmentation cases are marked with red (15 nm AuNP) and gray shaded area (5 nm AuNP).

**Table S1. Parameters in TTM model and MD simulation**

| Parameters | Value | Unit | Ref. |
|---|---|---|---|
| Thermal conductivity of electron $K_e$ | $K_e = \chi \frac{(\phi_e{}^2+0.16)^{5/4}(\phi_e{}^2+0.44)\phi_e}{(\phi_e{}^2+0.092)^{1/2}(\phi_e{}^2+0.16\phi_l)}$ | W/(m·K) | 1 |
| Normalized electron temperature, $\varphi_e$ | $\phi_e = T_e/T_f$ | 1 | 1 |
| Normalized electron temperature, $\varphi_l$ | $\phi_l = T_l/T_f$ | 1 | 1 |
| Fermi temperature, $T_f$ | 64200 | K | 1 |
| Thermal conductivity of lattice, $K_l$ | 2 | W/(m·K) | 2 |
| Thermal conductivity of water, $K_m$ | $K_m$ = -0.9003748+0.008387698×$T^1$-1.118205×$10^{-5}$×$T^2$ | W/(m·K) | 3 |
| Specific heat of electron, $C_e$ | From VSAP | J/(kg·K) | 4 |
| Specific heat of lattice, $C_l$ | 129 for solid, 163 for liquid | J/(kg·K) | 3 |
| Specific heat of water, $C_m$ | 4035.841+0.492312×$T^1$ for $T_m$<373K,<br>4219 for $T_m$>373K | J/(kg·K) | 3 |
| Density of gold, $\rho_{Au}$ | 19501.44-0.6933844×$T^1$-2.041944×$10^{-4}T^2$+4.297982×$10^{-8}T^3$ ~19300 for 86K<$T_l$<1338K<br>19033-1.4434×T for 1338K<$T_l$<3080K<br>14587.4 for $T_l$>3080K | kg/$m^3$ | 3 |
| Bulk melting temperature, $T_m^*$ | 1337 | K | 3 |
| Interface tension between liquid and solid gold, $\sigma_{sl}$ | 0.27 | N/m | 5 |
| Density of water, $\rho_m$ | 972.7584+0.2084×T^1-4.0×$10^{-4}$×$T^2$ for 273K<$T_m$<283K<br>345.28+5.749816×$T^1$-0.0157244×$T^2$+1.264375×$10^{-5}$×$T^3$ for 283K<$T_m$<373K<br>958 for $T_m$>373K | kg/$m^3$ | 3 |
| Electron-phonon coupling factor, $G_{e-l}$ | Calculated from VASP | W/($m^3$·K) | 4 |
| Interfacial thermal conductance, h | 105×$10^6$ | W/($m^2$·K) | 6 |
| Electron density of states, EDOS | Calculated from VASP | 1/($cm^3$·$eV^1$) | 4 |
| Chemical potential, μ | Calculated from VASP | J/kg | 4 |
| Boltzmann constant, $k_B$ | 1.38×$10^{-23}$ | $m^2$·kg/($s^2$·K) | 1 |
| Unit cell edge length of gold, $a_{fcc}$ | 4.08 | Å | 7 |
| Volume of AuNP, $V_{NP}$ | $V_{NP} = {}^4/_3\, \pi R_{NP}{}^3$ | $nm^3$ | 7 |

| | | | |
|---|---|---|---|
| Radius of AuNP, $R_{NP}$ | 5~100 | nm | |
| Wigner-Seitz radius, $r_{ws}$ | $1.65\times10^{-8}$ | cm | 7 |
| Surface tension of gold, $\sigma$ | 8900 for $T_l<1337$<br>$1150+0.14(T_l-1337)$ for $T_l>1337$ | dyne/cm | 8 |
| Elementary charge, e | $4.803204\times10^{-10}$ | stat C | 7 |
| Stretching stiffness, $K_s$ | 450.00 | kcal/(mol·Å$^2$) | 9-11 |
| Equilibrium bond length, $r_0$ | 0.9572 | Å | 9-11 |
| Bending stiffness, $K_\Theta$ | 55.00 | kcal/(mol·rad$^2$) | 9-11 |
| Equilibrium angle, $\Theta_0$ | 104.52 | degree | 9-11 |
| Partial atomic charge of O, $q_O$ | -0.83 | C | 9-11 |
| Partial atomic charge of H, $q_H$ | 0.415 | C | 9-11 |
| Well-depth of OO bond, $\varepsilon_{OO}$ | 0.102 | kcal/mol | 9-11 |
| LJ radius of OO bond, $\sigma_{OO}$ | 3.188 | Å | 9-11 |
| Well-depth of AuO bond, $\varepsilon_{AuO}$ | 0.59 | kcal/mol | 9-11 |
| LJ radius of AuO bond, $\sigma_{AuO}$ | 3.6 | Å | 9-11 |

Note: The bond energy ε and bond length σ for HH, OH, and AuH are set to 0.

**Table S2. Previous reported laser fluence of ps laser fragmentation for plasmonic nanoparticles.**

| Wavelength | Pulse duration | Material, diameter | Ligand | Fluence | Reference |
|---|---|---|---|---|---|
| 355 nm | 30 ps | Au, 25 nm | Citrate | 14 $mJ/cm^2$ | 12 |
| 532 nm | 10 ps | Au, 53 nm | No | 30 $mJ/cm^2$ | 13 |
| 400 nm | 0.15 ps | Au, 50 nm | Citrate | 6 $mJ/cm^2$ | 14 |
| 355 nm | 15 ps | Au, 59 nm | Citrate | 19.6 $mJ/cm^2$ | 15 |
| 355 nm | 18 ps | Ag, 65 nm | Citrate | 7.5 $mJ/cm^2$ | 16 |
| 1070 nm | 5 ps | Ag, 25 nm | No | 0.13 $mJ/cm^2$ | 17 |
| 532 nm | 10 ps | Au, 54 nm | No | >10 $mJ/cm^2$ | 18 |
| 532 nm | 28 ps | Au, 5~100 nm | Citrate | 1~2 $mJ/cm^2$ | This study |

**References (also included in paper):**


1. Chen, J. K.; Beraun, J. E.; Tham, C. L., Investigation of Thermal Response Caused by Pulse Laser Heating. *Numer. Heat Transfer, Part A* **2003,** *44* (7), 705-722.
2. Jain, A.; McGaughey, A. J. H., Thermal transport by phonons and electrons in aluminum, silver, and gold from first principles. *Physical Review B* **2016,** *93* (8), 081206.
3. COMSOL. Inc COMSOL Multiphysics® Reference Manual, version 5.3. www.comsol.com.
4. Lin, Z.; Zhigilei, L. V.; Celli, V., Electron-phonon coupling and electron heat capacity of metals under conditions of strong electron-phonon nonequilibrium. *Phys. Rev. B* **2008,** *77* (075133).
5. Font, F.; Myers, T. G., Spherically symmetric nanoparticle melting with a variable phase change temperature. *Journal of Nanoparticle Research* **2013,** *15* (12), 2086.
6. Plech, A.; Kotaidis, V.; Gresillon, S.; Dahmen, C.; von Plessen, G., Laser-Induced Heating and Melting of Gold Nanoparticles Studied by Time-Resolved X-Ray Scattering. *Phys. Rev. B* **2004,** *70*, 195423.
7. Giammanco, F.; Giorgetti, E.; Marsili, P.; Giusti, A., Experimental and Theoretical Analysis of Photofragmentation of Au Nanoparticles by Picosecond Laser Radiation. *J. Phys. Chem* **2010,** *114*, 3354-3363.
8. Aqra, F.; Ayyad, A., Theoretical temperature-dependence surface tension of pure liquid gold. *Mater. Lett.* **2011,** *65* (14), 2124-2126.
9. MacKerell, A. D.; Bashford, D.; Bellott, M.; Dunbrack, R. L.; Evanseck, J. D.; Field, M. J.; Fischer, S.; Gao, J.; Guo, H.; Ha, S.; Joseph-McCarthy, D.; Kuchnir, L.; Kuczera, K.; Lau, F. T. K.; Mattos, C.; Michnick, S.; Ngo, T.; Nguyen, D. T.; Prodhom, B.; Reiher, W. E.; Roux, B.; Schlenkrich, M.; Smith, J. C.; Stote, R.; Straub, J.; Watanabe, M.; Wiórkiewicz-Kuczera, J.; Yin, D.; Karplus, M., All-Atom Empirical Potential for Molecular Modeling and Dynamics Studies of Proteins. *The Journal of Physical Chemistry B* **1998,** *102* (18), 3586-3616.
10. Jorgensen, W. L.; Chandrasekhar, J.; Madura, J. D.; Impey, R. W.; Klein, M. L., Comparison of simple potential functions for simulating liquid water. *The Journal of Chemical Physics* **1983,** *79* (2), 926-935.
11. Price, D. J.; III, C. L. B., A modified TIP3P water potential for simulation with Ewald summation. *The Journal of Chemical Physics* **2004,** *121* (20), 10096-10103.

12. Inasawa, S.; Sugiyama, M.; Yamaguchi, Y., Bimodal Size Distribution of Gold Nanoparticles under Picosecond Laser Pulses. *The Journal of Physical Chemistry B* **2005,** *109* (19), 9404-9410.
13. Ziefuß, A. R.; Reichenberger, S.; Rehbock, C.; Chakraborty, I.; Gharib, M.; Parak, W. J.; Barcikowski, S., Laser Fragmentation of Colloidal Gold Nanoparticles with High-Intensity Nanosecond Pulses is Driven by a Single-Step Fragmentation Mechanism with a Defined Educt Particle-Size Threshold. *The Journal of Physical Chemistry C* **2018,** *122* (38), 22125-22136.
14. Werner, D.; Furube, A.; Okamoto, T.; Hashimoto, S., Femtosecond Laser-Induced Size Reduction of Aqueous Gold Nanoparticles: In Situ and Pump−Probe Spectroscopy Investigations Revealing Coulomb Explosion. *The Journal of Physical Chemistry C* **2011,** *115* (17), 8503-8512.
15. Hashimoto, S.; Katayama, T.; Setoura, K.; Strasser, M.; Uwada, T.; Miyasaka, H., Laser-driven phase transitions in aqueous colloidal gold nanoparticles under high pressure: picosecond pump-probe study. *Phys. Chem. Chem. Phys.* **2016,** *18* (6), 4994-5004.
16. Kamat, P. V.; Flumiani, M.; Hartland, G. V., Picosecond Dynamics of Silver Nanoclusters. Photoejection of Electrons and Fragmentation. *The Journal of Physical Chemistry B* **1998,** *102* (17), 3123-3128.
17. Jeon, J.-W.; Yoon, S.; Choi, H. W.; Kim, J.; Farson, D.; Cho, S.-H., The Effect of Laser Pulse Widths on Laser—Ag Nanoparticle Interaction: Femto- to Nanosecond Lasers. *Applied Sciences* **2018,** *8* (1), 112.
18. Ziefuss, A. R.; Reich, S.; Reichenberger, S.; Levantino, M.; Plech, A., *In Situ* Structural Kinetics of Picosecond Laser-Induced Heating and Fragmentation of Colloidal Gold Spheres. *Phys. Chem. Chem. Phys.* **2020,** *22*, 4993-5001.
